# Label-free multiphoton imaging reveals volumetric shifts across development in sensory-related brain regions of a miniature transparent vertebrate

**DOI:** 10.1101/2024.07.18.604134

**Authors:** Rose L. Tatarsky, Najva Akbari, Ke Wang, Chris Xu, Andrew H. Bass

## Abstract

Animals integrate information from different sensory modalities as they mature and perform increasingly complex behaviors. This may parallel differential investment in specific brain regions depending on the demands of changing sensory inputs. To investigate developmental changes in the volume of canonical sensory integration brain regions, we used third harmonic generation imaging for morphometric analysis of forebrain and midbrain regions from 5 to 90 days post fertilization (dpf) in *Danionella dracula*, a transparent, miniature teleost fish whose brain is optically accessible throughout its lifespan. Relative to whole brain volume, increased volume or investment in telencephalon, a higher order sensory integration center, and torus longitudinalis (TL), a midbrain visuomotor integration center, is relatively consistent from 5 to 30 dpf, until it increases at 60 dpf, followed by another increase at 90 dpf, as animals reach adulthood. In contrast, investment in midbrain optic tectum (TeO), a retinal-recipient target, progressively decreases from 30-90 dpf, whereas investment is relatively consistent across all stages for the midbrain torus semicircularis (TS), a secondary auditory and mechanosensory lateral line center, and the olfactory bulb (OB), a direct target of the olfactory epithelium. In sum, increased investment in higher order integration centers (telencephalon, TL) occurs as juveniles reach adulthood and exhibit more complex cognitive tasks, whereas investment in modality-dominant regions occurs in earlier stages (TeO) or is relatively consistent across development (TS, OB). Complete optical access throughout *Danionella*’s lifespan provides a unique opportunity to investigate how changing brain structure over development correlates with changes in connectivity, microcircuitry, or behavior.

## INTRODUCTION

Behavioral capabilities change over the course of development, in part, depending on age-related changes in the size and connectivity of sensory systems (e.g., Brandstätter and Kotrschal, 1990; Nixdorf-Bergweiler & von B. und Halbach, 2004; Gonda, Herczeg, and Merilä, 2009; Hinaux et al., 2016; Yi et al., 2022). As developing brain tissue is energetically expensive (Niven & Laughlin, 2008; O’Donnell et al., 2011; Kotrschal et al., 2013), differential growth, or investment, in brain regions may vary according to the demands of certain sensory inputs (Barton et al., 1995; Moran, Softley & Warrant, 2015; Stöckl et al., 2016; Sheehan et al., 2019; Keesey et al., 2020). The diversity of environmental niches shaping sensory specialization and behavioral repertoires in fish species has offered significant insights into how relative investment in different brain regions changes from larval stages to adulthood when they exhibit more complex cognitive tasks, such as courtship and aggression (e.g., Toyoda and Uematsu, 1994; Huber et al., 1997; Iribarne and Castello, 2014, Jaggard et al., 2020; Axelrod et al., 2018; Yi et al., 2022). Here, we used third harmonic generation (THG), multiphoton microscopy to map the age-related growth of sensory and sensory-related integration centers in *Danionella dracula*, a miniature, transparent cyprinid fish that along with other *Danionella* species has emerged as a new model for investigating the neural basis of adult social behavior (e.g., Schulze et al., 2018; Rajan et al., 2022; Tatarsky et al., 2022; Lee & Briggman, 2023; Zada et al., 2024).

Research into differences in brain region size and investment across development has traditionally relied on methods such as postmortem tissue sectioning and immunohistochemistry, as well as computed tomography (CT) for three dimensional (3D) visualization (Brandstätter and Kotrschal, 1990; Iribarne and Castelló, 2014; Yi et al., 2022). In small, transparent animals like larval zebrafish (*Danio rerio*) and larval cavefish (*Astyanax mexicanus*), intact whole-brain imaging approaches have gained prominence, enabling the generation of detailed three dimensional brain atlases and revealing anatomical variations between species populations (Ullmann et al., 2010; Kunst et al., 2019; Jaggard et al., 2020). However, studies using postmortem tissue prepared with traditional fixation methods may introduce artifacts such as the loss of extracellular space, potentially affecting morphometric analysis of different brain regions (Pallotto et al., 2015). Fortunately, recent advancements in microscopy have pushed the boundaries of optical accessibility, facilitating the imaging of live, intact animal brains (e.g., McArthur, Chow, Fetcho 2020; Chow et al., 2020; Akbari et al., 2022; Lam, 2022; Akbari et al., 2024). We built upon these advancements and imaged the brain of live *D. dracula* in different developmental stages with THG microscopy, a label-free multiphoton technique that detects tissue heterogeneities with micrometer resolution, as strong THG signal is generated by differences in nonlinear refractive index at interfaces such as those between water and lipids (e.g., Farrar et al., 2011; Weigelin, Bakker, and Friedl, 2016; Ahn et al., 2020). Previously, THG microscopy has been used for *in vivo* imaging of myelination and blood vasculature in the brain without the need for any exogenous label (Farrar et al., 2011; Ahn et al., 2020; Chow et al., 2020).

*Danionella dracula* and other *Danionella* species are miniature, paedomorphic vertebrates that remain transparent through their entire lifespan (Britz et al., 2009; Britz and Conway, 2016; Conway, Kubicek and Britz; 2021) (Fig. 1A, B). Their small size, lack of ossified dorsal skull, and lifelong transparency offer significant advantages for brain imaging using optical microscopy, including THG (Schulze et al., 2018; Chow et al., 2020; Britz, Conway, and Rüber, 2021; Akbari et al., 2022; Akbari et al., 2024). This provides a distinct opportunity for non-invasive investigation of entire brain regions and their subdivisions from early development to adulthood, and allows for future comparisons of neurodevelopmental dynamics between paedomorphic species and other teleost fish species (Britz & Conway, 2016; Conway, Kubicek, and Britz, 2021).

**Figure 1.**
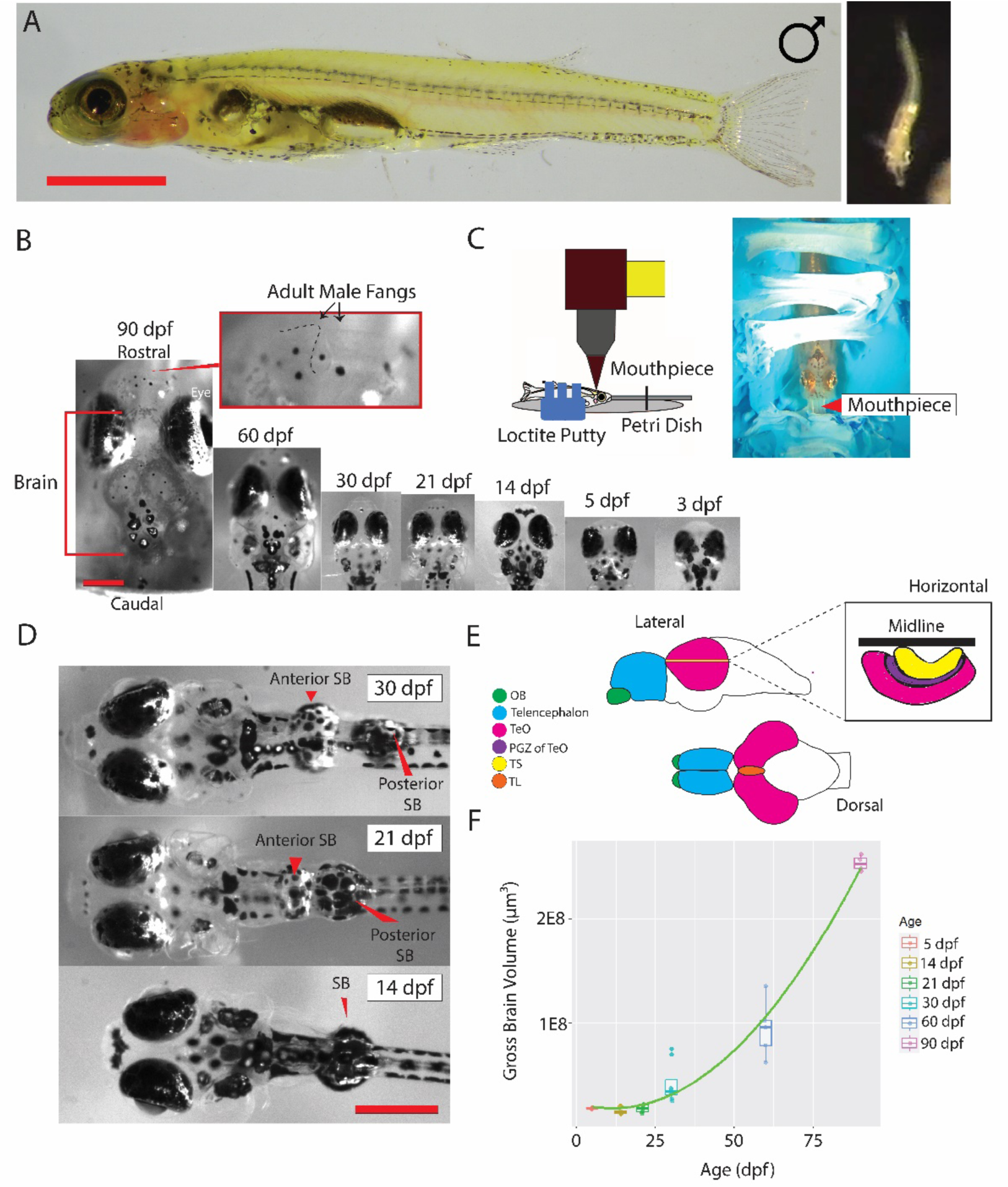
Third harmonic generation (THG) brain imaging in groups of *Danionella dracula* 3 to 90 dpf. **A**. Left: Side view of a male. Scale bar indicates 3mm. Right: Head-on view of male extending its hypertrophied lower jaw during mid-lunge at another male. **B**. Dorsal views of fish imaged from 90-3 dpf (left to right). Scale bar left indicates 500 μm for all images. Top insert at 90 dpf is zoom-in on fangs (indicated by arrows; dashed outline on left fang), which are first apparent in 90 dpf males along with hypertrophied lower jaw. **C**. Schematic of imaging apparatus (see Materials and Methods). Left: profile drawing of animal placement under objective. Right: dorsal view of adult placed in putty holder with perfusion mouthpiece. Fish imaged less than 60 dpf were imaged using an alternative setup than pictured (see Materials and Methods, Fish stabilization for imaging). **D.** 30, 21, and 14 dpf example fish depicting changes in swim bladder (SB) separation into anterior and posterior chambers. Scale bar indicates 500 μm for all images. **E**. Schematic of representative *D. dracula* brain; illustrations based on pictures of dissected whole brains. Brain regions of morphometric analysis indicated in lateral and dorsal views; green, OB (olfactory bulb); blue, telencephalon; orange, TL (torus longitudinalis); pink, TeO (optic tectum). Insert indicates horizontal slice in midbrain depicting TeO (pink), periventricular gray zone (PGZ) of TeO (purple, included in TeO measurement), depicted for indication of boundary between TeO and TS (yellow, torus semicircularis). **F**. Scatterplot of age in dpf of animals in this study and gross brain volume in μm^3^. Boxplots indicate median, upper and lower quartiles and 1.5× interquartile range for each Age. Gross brain volume calculation is described in Table 2 and Materials and Methods. Different ages depicted by different colored dots as shown in legend: red, 5 dpf; yellow, 14 dpf; green, 21 dpf; teal, 30 dpf; blue, 60 dpf; purple, 90 dpf. As is described in Materials and Methods, we performed the best fitting quadratic regression (F_2,30=_120.8, R^2^= 0.8821, p<0.001) on log-transformed data to normalize the residuals. The green line displayed here represents the quadratic regression line based on the original (non-transformed) data, for best visual clarity and understanding.

Here, we used THG imaging and morphometric analysis in live *D. dracula* from early developmental stages (3 days post-fertilization, dpf) until adulthood (90 dpf) to investigate changes in the relative investment in brain regions deemed especially relevant to sensory mechanisms coupled to acoustic and postural displays in adults (see Tatarsky et al., 2022). The selected regions were also chosen, in part, because of the ease of recognizing defined boundaries using THG across several developmental stages (see Materials and Methods). This included three primary or secondary targets of the sensory periphery that we refer to as modality-dominant: the forebrain olfactory bulb (OB), a primary target of the olfactory epithelium; the midbrain optic tectum (TeO), a primary target of retinal ganglion cells; the midbrain torus semicircularis (TS), a secondary target of mechanosensory auditory and, in aquatic vertebrates, lateral line hair cell epithelia (Striedter, 2005; Nieuwenhuys, ten Donkelaar and Nicholson, 1998). Additionally, two higher order sensory integration centers were studied: the midbrain torus longitudinalis (TL), which has strong connections to the TeO, receives inputs from cerebellum-related nuclei of the pretectum, and is largely implicated in the integration of visual information (DeMarco e al., 2021; Tesmer et al., 2022), and the telencephalon, a multisensory integration site (e.g., Lopes Corrêa, Grant, and Hoffmann, 1998; Prechtl et al., 1998; Folgueira, Anadón, and Yáñez, 2004; Striedter and Northcutt, 2022) that is further implicated across vertebrate taxa, including teleost fish, in the executive control of complex behaviors such as social interactions, learning and memory (Davis & Northcutt, 1983; Shinozuka and Watanabe, 2004; Teles et al., 2015; Giassi et al., 2012; Stednitz et al., 2018).

Species-specific patterns in relative investment in different brain regions has been reported to emerge during late larval and juvenile periods in various species of cyprinids (Brandstätter and Kotrschal, 1989, 1990). Therefore, we hypothesized that brain regions known to be involved in distinct sensory functions would grow allometrically compared to the rest of the brain as *D. dracula* larvae become juveniles, and especially in fish ∼60-90 dpf (2-3 months) of age, the period when juveniles transform into adults and exhibit sex-typical morphology (e.g., male fangs and hypertrophied lower jaw) and social behaviors, including aggression and courtship (see Tatarsky et al., 2022).

## MATERIALS AND METHODS

### Fish Care and Egg Collection

Animal husbandry and rearing was conducted as is previously described (Tatarsky et. al., 2022), and all procedures were approved by the Institutional Animal Care and Use Committee of Cornell University. Briefly, adults were bred in a laboratory colony at Cornell. Eggs were collected from breeding tanks by removing the nest, unrolling the filtration cartridge, and gently moving the clusters of eggs to acrylic cylinders (10 cm diameter) with a mesh bottom that rested within a 50-gal aquarium. Larval fish were fed AP100 Dry Larval Diet (Zeigler Bros, Inc.) twice a day in these smaller cylinders, supplemented with L-type rotifers (*Brachionus plicatilis)*.

### Fish Rearing and Collection for Imaging

For imaging, 3 and 5 dpf larvae were removed gently from their respective smaller cylinders by scooping them out of the water using a small plastic cup and then transferred into a petri dish containing 50 ml of system water (see Table 1 for sample size and standard length, SL, for each age group imaged). For stages after 10 dpf, fish who had hatched from eggs which were all laid on the same day were moved from initial acrylic cylinders to three separate 10-gal tanks (n∼50 larvae in each tank) where they were fed appropriate diet for age and raised in colony settings described previously. All three tanks were maintained for proper water chemistry and did not experience fluctuations in parameters, and there were no differences in chemistry between the tanks. Fish were removed from the tank for imaging using a plastic cup at the following ages: 14, 21, 30, 60 and 90 dpf, which is when these fish become adults as determined without the aide of microscopy by the presence of eggs in the abdomen of females and the hypertrophied jaw and fangs in males (Britz, Conway and Rüber, 2009; Britz and Conway, 2016; Tatarsky et. al., 2022). These adult characteristics also enabled the fish to be reliably sexed at 90 dpf, which was not possible given the absence of these traits in earlier stages. We imaged fish reared exclusively in tanks containing individuals hatched on the same day to control for age, and fish were selected to be similar in size for each age group imaged and to be representative of typical size at these ages, avoiding the selection of especially small or large animals to represent the age group.

**Table 1:** Standard Length Ranges of Imaged Age Groups

| Stage | 3 dpf | 5 dpf | 14 dpf | 21 dpf | 30 dpf | 60 dpf | 90 dpf |
| --- | --- | --- | --- | --- | --- | --- | --- |
| Number of animals imaged (n) | 5 | 5 | 5 | 6 | 8 | 5 | 4 |
| Sex | See above on inability to sex fish before 90 dpf | - | - | - | - | - | 3 male, 1 female |
| Standard Length Range (mm) | 3.1-3.8 | 3.3-3.8 | 3.5-4.0 | 3.2-3.8 | 3.7-5.2 | 6.8-8.1 | 11.9-13.9 |

### Fish Stabilization for Imaging

Before imaging, *D. dracula* were anesthetized in different dilutions of a benzocaine solution depending on stage of development as follows: 0.0015% for 3-21 dpf, 0.002% for 30 dpf, 0.0025% for 60-90 dpf. Fish aged 3-30 dpf were mounted dorsal side up in 1.5% low-melting temperature agarose (Sigma-Aldrich) on a glass bottom dish glued to the bottom of a petri dish chamber. In 14-30 dpf fish, after the agarose gelled, agarose covering the mouth and gills was carefully removed with a needle without touching the fish, and mouthpiece tubing was secured anterior to the mouth to perfuse well-oxygenated temperature-controlled fish system water (2 L reservoir heated with a Top Fin Betta Aquarium Heater set to 25 °C) containing benzocaine solution of 0.006% and 0.008% for 5-21 and 30 dpf, respectively (Fig. 1C). These doses kept the fish stationary during neuroimaging, and the reduction in anesthesia perfused during imaging was so the fish will be anesthetized and stable during the experiment but easily revived following the procedure (Schulze et al., 2018). Perfusion occurred at a rate of 700 μl min^−1^ in 14-30 dpf fish, providing a slow steady drop rate of water over the gills. Another tube pulled water out of the dish at the same rate as perfusion to keep the water recirculating and to prevent the dish from overflowing. Fish that were 60-90 dpf were stabilized by positioning them on a mountable putty (Loctite 1865809) and perfused through the mouth and over the gills at a rate of 1 ml min^−1^. Each fish’s mouthpiece was custom made depending on its mouth size and delivered system water containing a 0.00185% benzocaine solution. All fish imaged and analyzed survived the procedure; survival was confirmed by observing blood flow in the brain throughout imaging and by reviving the fish to swim in a petri dish containing system water after imaging. Fish acclimated from imaging in the recovery petri dish for ∼30 mins before being placed back in the fish facility, into a separate colony tank from unimaged fish.

### Image Acquisition

Third harmonic generation images were obtained using a commercially available multiphoton microscope (Bergamo II system B242, Thorlabs Inc.) with a high numerical aperture objective lens (XLPLN25XWMP2, Olympus, NA 1.05). The laser source was a non-collinear optical parametric amplifier (NOPA, Spectra Physics) pumped by an amplifier (Spirit 1030-70, Spectra Physics). To control the excitation power, a half waveplate and a polarization beam splitter were used. A two-prism (SF11 glass) compressor was used to compensate for the normal dispersion of the optics of the light source and the microscope, including the objective. A laser wavelength of 1280 nm with a repetition rate of 333 kHz was used for signal generation. The full width at half maximum (FWHM) pulse width was measured to 50 fs at 1280 nm, assuming a sech^2^ pulse intensity profile.

Images were collected at approximately 1 frame per second over a field-of-view (FOV) of 539 µm by 539 µm with 512 by 512 pixels for all images in 60-90 dpf fish, and for images of the midbrain and hindbrain of 5-30 dpf fish. The FOV for images of the telencephalon in 5-30 dpf fish was 303.16 µm by 303.16 µm with 512 by 512 pixels to avoid inclusion of the eyes in images, which produce a large amount of background signal. The FOV for all images in 3 dpf fish was 432.52 µm by 432.52 µm with 512 by 512 pixels. To reduce imaging time and facilitate health and survival, the midbrain and hindbrain were only imaged unilaterally in their respective stacks rather than tiling across the entirety of these regions of the brain. There was no apparent difference in the size of one half of the brain compared to the other. The telencephalon was always imaged in entirety due to its size fitting entirely in one FOV. Images were acquired in 1.5-5 µm steps in the Z direction based on size, where smaller animals were imaged with smaller step sizes to better resolve the smaller regions of interest (ROIs) compared to imaged larger animals.

### Brain Region Segmentation

All segmentation of individual brain regions was performed using Imaris Bitplane software (Oxford Instruments). Multiphoton imaging stacks were imported into Fiji/ImageJ for scaling and then scaled tifs were imported to Imaris (9.5.1). The color of the imported stacks was made gray, and histograms were adjusted for best visualization of boundaries between different brain regions. Gaussian filters were applied to images before analysis for best image contrast. Focal brain regions were manually segmented by tracing the outline of a focal brain region on individual slices in the horizontal plane (XY) using the boundary definitions described in Fig. 2 and Table 2. Boundaries were confirmed by outlining slices in the other planes of view (XZ, YZ). Brain atlases of larval and adult zebrafish were used to guide segmentation and to create boundary definitions (Wullimann, Rupp, and Reichert, 1996; Kunst et al., 2019). The regions segmented were the telencephalon (bilateral), TeO (unilateral), TS (unilateral; not sampled at 5 dpf, see below), TL (midline; not sampled at 5, 14, and 21 dpf, see below) and OB (bilateral; not sampled at 90 dpf, see below). The telencephalon measurement included the anterior portion of the preoptic area (POA). The volumes of each TeO and TS were doubled for analyses, as these structures appeared to be bilaterally symmetrical. We did not measure TL for animals aged 5-21 dpf because TL could not be differentiated from the midbrain with THG at those ages. Similarly, TS could not be differentiated in animals aged 5 dpf so we did not measure TS at that age. The OB for 90 dpf animals only included one animal as the imaging field of view cut off most of both OBs in three of four 90 dpf animals. For each region, ∼40-50 slices in the horizontal plane were traced with uniform spacing across all ages to account for differences in brain region size in the different ages and to have similar resolution to each other.

**Figure 2.**
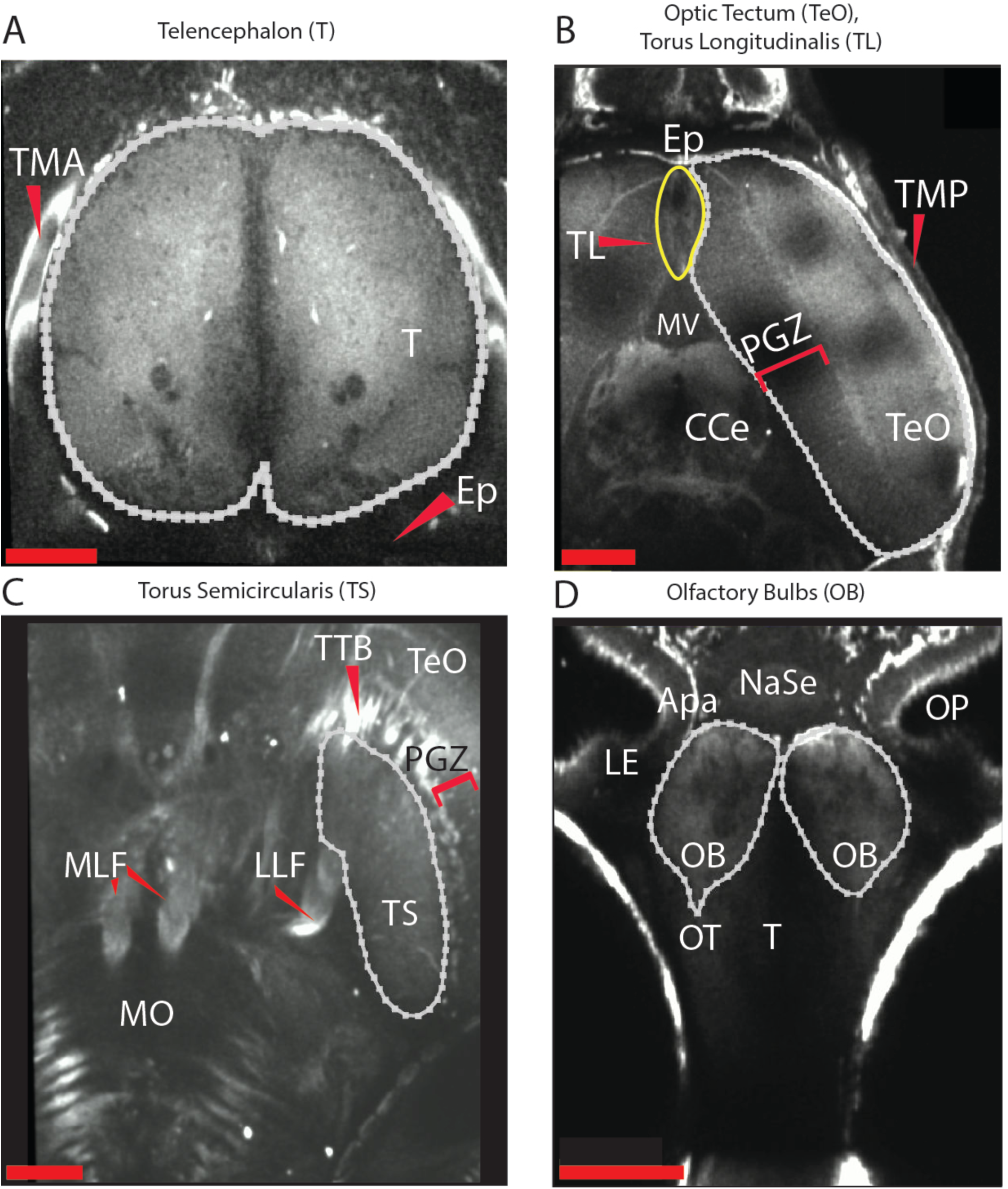
Brain Region Segmentation using third harmonic generation (THG) microscopy. Images from representative horizontal slices of 60 and 90 dpf fish. Strong THG signal is in white, and dense regions containing neuronal cell bodies (somata) appear as cell shadows or darker regions (also see Akbari et al., 2024). Top = anterior, rostral end of fish; bottom = posterior, caudal end of fish; left = left; right = right. All boundaries and abbreviations defined in Table 2. Red scale bars indicate 100 μm. A) Image from imaging stack containing 90 dpf telencephalon (T) (white outline). B) Image from stack containing 60 dpf TeO (white outline) and TL (yellow outline), other midbrain structures, and cerebellum region. Corpus cerebelli (CCe) traced and included in the gross brain volume measurement described in Table 2. C) Image from stack containing 90 dpf midbrain and hindbrain regions which contains TS (white outline). D) Image from stack containing 60 dpf T and OB (white outline), which lie ventral and anterior to T. Slice here taken from section of OB ventral to T, thus the bottom of T is only beginning to enter view.

**Table 2.**
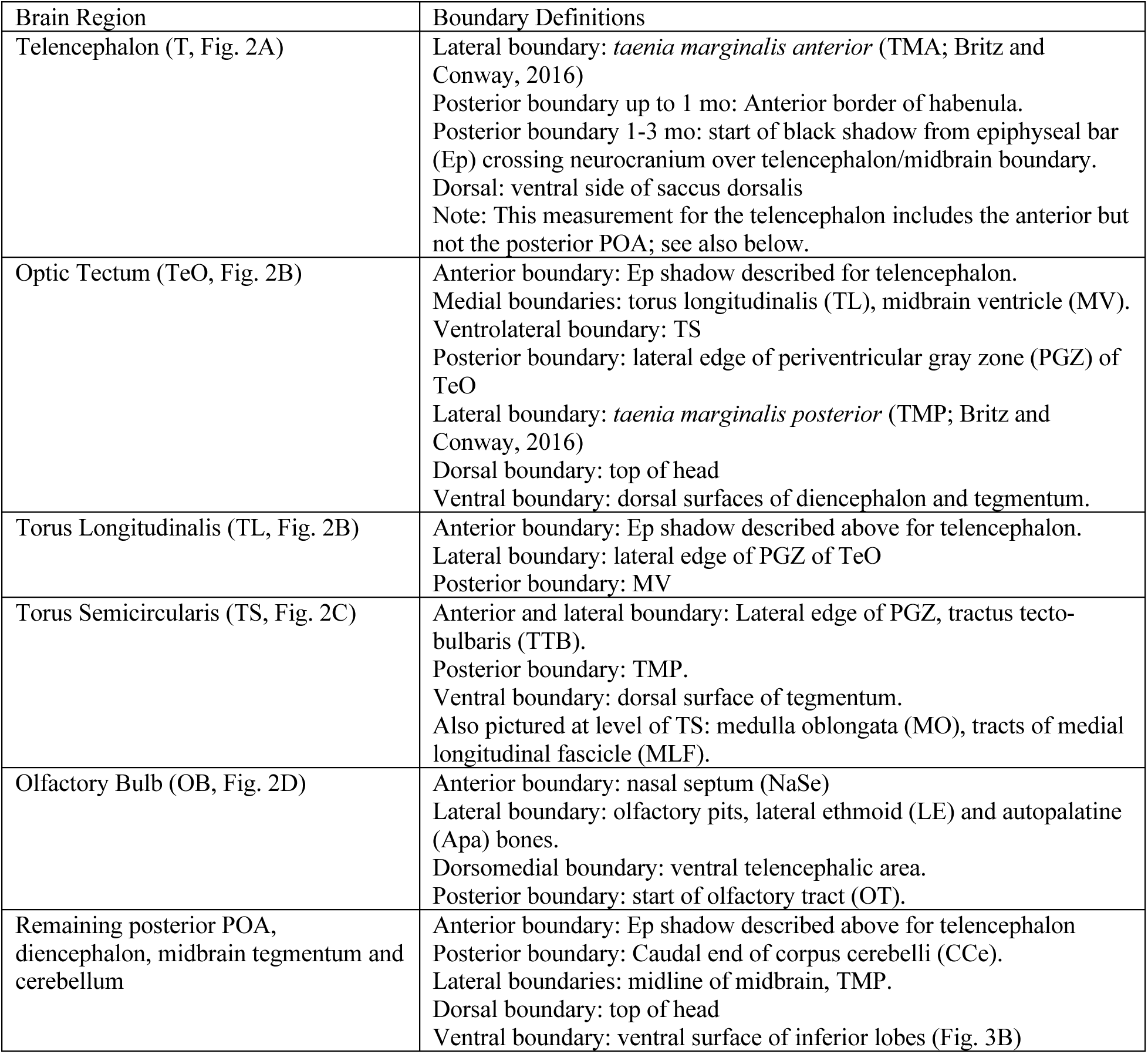
Brain Region Segmentation Boundary Definitions

We also segmented a broader region containing areas difficult to differentiate with clear boundaries, but that we wanted to include when calculating “gross brain volume” (see below for calculation) that could be demarcated in all age groups and would allow for comparisons in relative size differences of the TeO, TS, TL, OB and telencephalon (see Methods: Morphometric and Statistical Analyses). This measurement included the cerebellum, posterior preoptic area and diencephalon, and the rest of the remaining midbrain tegmentum; it excluded the already measured TeO, TL, and TS. The measurement spanned from the telencephalon-midbrain boundary to the caudal end of the corpus cerebelli along the midline (see Table 2 for boundary definitions for remaining midbrain tegmentum and cerebellum).

### Morphometric and Statistical Analyses

Morphometric image analysis was performed using the software Imaris (BITPLANE, Oxford Instruments). Following segmentation, brain regions were created into 3D surface objects, or computer-generated representations of the brain regions based on the individual slices traced, in Imaris. This surface object is an artificially solid object which allows the visualization and quantification of the volume of the traced brain region (Imaris 9.2 Reference Manual, BITPLANE, Oxford Instruments). Volume statistics for a brain region were automatically quantified in Imaris Bitplane from the areas of the traced slices (the sum of the triangle surfaces for a traced region) and the distance between traced slices (Imaris 9.2 Reference Manual, BITPLANE, Oxford Instruments).

To standardize for differences in brain size across ages, we converted absolute volumes to relative volumes compared to the gross brain volume. This was done by dividing each absolute volume by the gross brain volume, which was calculated by summing the volumes of both telencephali, 2x the TeO volume, 2x the TS volume, the volume of the midline TL, and the volume of the remaining posterior POA, diencephalon, midbrain tegmentum (MT; midbrain without the TeO, TL and TS) and cerebellum. The cerebellum provided an easily demarcated caudal landmark in THG images for the broad extent of the whole brain studied here (see Table 2). The volumes of OB were not included in the gross brain volumes because they could not be entirely measured in three of four 90 dpf animals. However, we do note that the olfactory bulbs can be distinctly differentiated from the telencephalon in every animal. Relative volumes were then converted into percentages of the gross brain volume.

### Statistical analyses

Statistical analyses were performed in R version 4.1.2 (R Core Team, 2021). To determine the best fitting regression for gross brain volume by age (dpf), we compared linear, quadratic and cubic models of the data. As gross brain volume data were not normally distributed, we used log (1+*x*) transformation to normalize the data. Shapiro–Wilk tests confirmed that transformation resulted in normally distributed residuals. In Figure 1F, we present the quadratic regression line using the original (non-transformed) data to enhance visual clarity and comprehension.

Statistical differences in brain region size were determined by performing a one-way ANOVA for each brain region to examine the effect of age group and region on the normalized brain percentage of the region. In the case of significant main effects or interactions, we followed these measures with *post hoc* tests using the *lsmeans* package (Lenth, 2016). We corrected for multiple comparisons using the Benjamini-Hochberg method.

We also used linear mixed models to examine the effect of brain region and body length on absolute brain volumes for two pooled age groups (animals aged 30 dpf and younger, and older animals of 60 and 90 dpf). We ran a separate model for each comparison, comparing changes in absolute brain volume for each of the different brain regions (TeO, TL, TS, OB) in relation to the telencephalon. The coefficients of the predictor variables (Brain Region and Standard Length) were used to derive the slopes in Table 11 that quantified the relationship between body length and brain region concerning absolute brain volume.

## RESULTS

To compare the size of different brain regions relevant to sensation across different ages of *D. dracula,* we performed THG imaging in groups of fish 3-90 dpf. We considered adults to be fish with secondary sex characteristics visible to the naked eye, namely the hypertrophied lower jaw and fangs in males and the presence of eggs in females (right, Fig. 1A). Larvae, juveniles, and adults are transparent and thus their brains amenable to whole-brain imaging in intact animals *in vivo* throughout the entirety of development (Fig. 1B, D, E). The imaging holder for 60 and 90 dpf fish is shown in Fig. 1C; younger ages were embedded in agarose (see Materials and Methods). The size of individual brain regions within each age group was quantified by manually segmenting and quantifying the size of the telencephalon, TeO, TS, TL, and OB, into distinct anatomical regions for all brains imaged (Fig. 2). This was done in accordance with previous descriptions in zebrafish (Wullimann, Rupp, and Reichert, 1996; Kunst et al., 2019). We controlled for age by imaging fish reared exclusively in tanks containing individuals hatched on the same day; those selected for imaging were chosen to be as representative and similar in size to other fish of the same age as possible, to address potential concerns regarding the conflation of age and standard length.

### Somatic and behavioral characters in 3-90 dpf *D. dracula*

At 3 dpf, *D. dracula* larvae have an intact yolk and are largely immotile. They can swim for short periods following disturbance of their water, but primarily they lie on the bottom of the holding tube. Their swim bladder was not inflated. At 5 dpf, the larvae’s swim bladder was inflated, and they swim at the top of the water column. The animals in these stages were very similar in size (see Table 1 in Materials and Methods for all SL information and number of animals imaged), but distinct in appearance due to differences in behavior, extent of swim bladder inflation, and loss of yolk in 5 dpf animals. Fish were reared in larger community tanks after 10 dpf rather than the larvae rearing tubes. The fish change their behavior from 14-30 dpf, initially swimming at the top of the water column to swimming throughout the tank at all levels. We observed from 14-30 dpf that the anterior portion of the swim bladder separates from the posterior portion (Fig. 1D). The fish also appear to begin to increase in size during this period (Table 1, Fig. 1E).

The mean standard length of animals aged 60 dpf that we imaged was approximately two times larger than the mean size of fish aged 30 dpf and younger, and animals aged 90 dpf were about 3.3 times larger (Table 1). At 60 dpf, juveniles possess extended fins and fin rays, resembling adult fish except for their smaller size and lack of secondary sex characteristics. Thus, we were not able to determine the sex of 60 dpf fish. *Danionella dracula* become adults around 90 dpf; this is readily recognizable by the presence of eggs in the abdomen of females and the hypertrophied jaw and fangs in males (right, Fig. 1A) (Tatarsky et. al., 2022, Britz, Conway and Rüber, 2009, Britz and Conway, 2016). We do not make comparisons between the brain regions of the different sexes at this time because we only imaged one female animal.

### Third Harmonic Generation (THG) Imaging

THG signal detection and contrast between brain region boundaries increases in quality based on age. Given this, we found that the boundaries between brain regions in 3 dpf animals were not possible to segment because the brains appeared largely homogenous in THG imaging. By at least 5 dpf, the brain regions were more robustly differentiated and thus reliably measured for all age groups, except for the TS which could not be differentiated in 5 dpf animals. Key features of the telencephalon can be seen in THG, including shadows of the the lateral forebrain bundle and anterior commissure (LFB, AC, respectively; Fig. 3A, B). The posterior end of the area dorsalis of the telencephalon appears to be segmented into discrete lobes (see blue outlines in Fig. 3B). The OBs are differentiated from the telencephalon, so we separate the quantification of these structures. Some commissures such as the posterior commissure (PC, Fig. 3C) can be seen in bright white, indicating strong THG signal. The AC, however, appears dark as a shadow (Fig. 3B) compared to other commissures. Dense regions containing cell bodies appear as cell shadows or darker regions, for example, the closely packed cells of the periventricular gray zone of the TeO (PGZ, Fig. 3C; also see Akbari et al., 2024).

**Figure 3.**
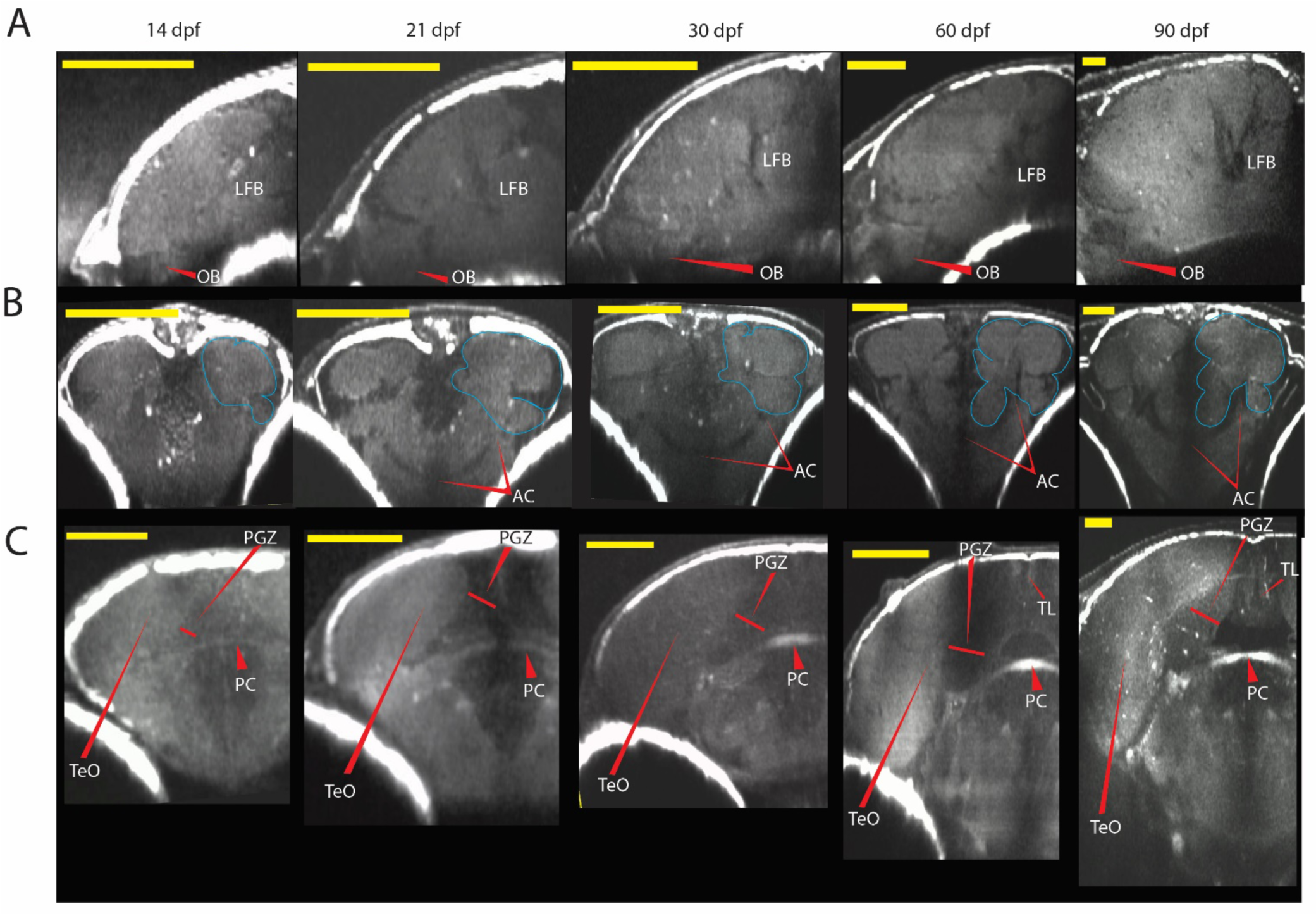
Representative third harmonic generation (THG) image slices for age groups 14-90 dpf. All scale bars indicate 100 μm. Images are not on the same scale because of the different sizes of brain regions across the five ages shown. Images were expanded so that labeled features could be easily visualized across the different ages. A. Sagittal slice taken at midline of telencephalon in 14-90 dpf animals. Red arrow indicates olfactory bulb (OB). LFB indicates tracts of lateral forebrain bundle. Left = anterior, right = posterior, top = dorsal, bottom = ventral. B. Coronal view from rostral perspective of posterior telencephalon in 14-90 dpf animals. Red pointers indicate extent of anterior commissure (AC) shadow, with one arrow pointing to the AC at the midline and the other to its extent on the lateral side of the brain. Blue lines on one side indicate the discrete lobes of the area dorsalis of telencephalon. Right = left, left = right, top = dorsal, bottom = ventral. C. Coronal view from rostral perspective of anterior midbrain optic tectum (TeO) and remaining midbrain in 14-90 dpf animals. Red arrows indicate TeO and periventricular gray zone of TeO (PGZ). PC indicates posterior commissure. Right = left, left = right, top = dorsal, bottom = ventral.

### Morphometric Analysis of Sensory Integration and Sensation Brain Regions

We used THG imaging in fish of various age groups to analyze and segment brain regions associated with sensory integration and sensation, including the telencephalon, TeO, OB, TL, and the TS (see Fig. 2). All examined regions increased in absolute volume as animals matured (Table 3). To determine the best fitting regression for changes in absolute gross brain volume by age (dpf), we compared linear, quadratic and cubic models. Quadratic regression provided the best fit, explaining a significant proportion of the variance compared to linear regression (linear: F_2,30 =_ 71.47, R^2^ = 0.6877, p<0.001; quadratic: F_1,31 =_ 120.8, R^2^ = 0.8821, p<0.001; Fig. 1F). Although the cubic regression showed marginal improvement over the quadratic regression (F_3,29 =_ 84.58, R^2^ = 0.8947, p<0.001), we considered the quadratic model to be the more parsimonious choice for modeling this data. Therefore, the relationship between gross brain volume and age best follows a nonlinear pattern, characterized by minimal changes in gross brain volume up to 30 dpf, followed by a substantial increase in later ages. This increase in the size of the brain coincides with the considerably larger changes in body size of animals aged 60 dpf and 90 dpf compared to younger ages (Table 1).

**Table 3.**
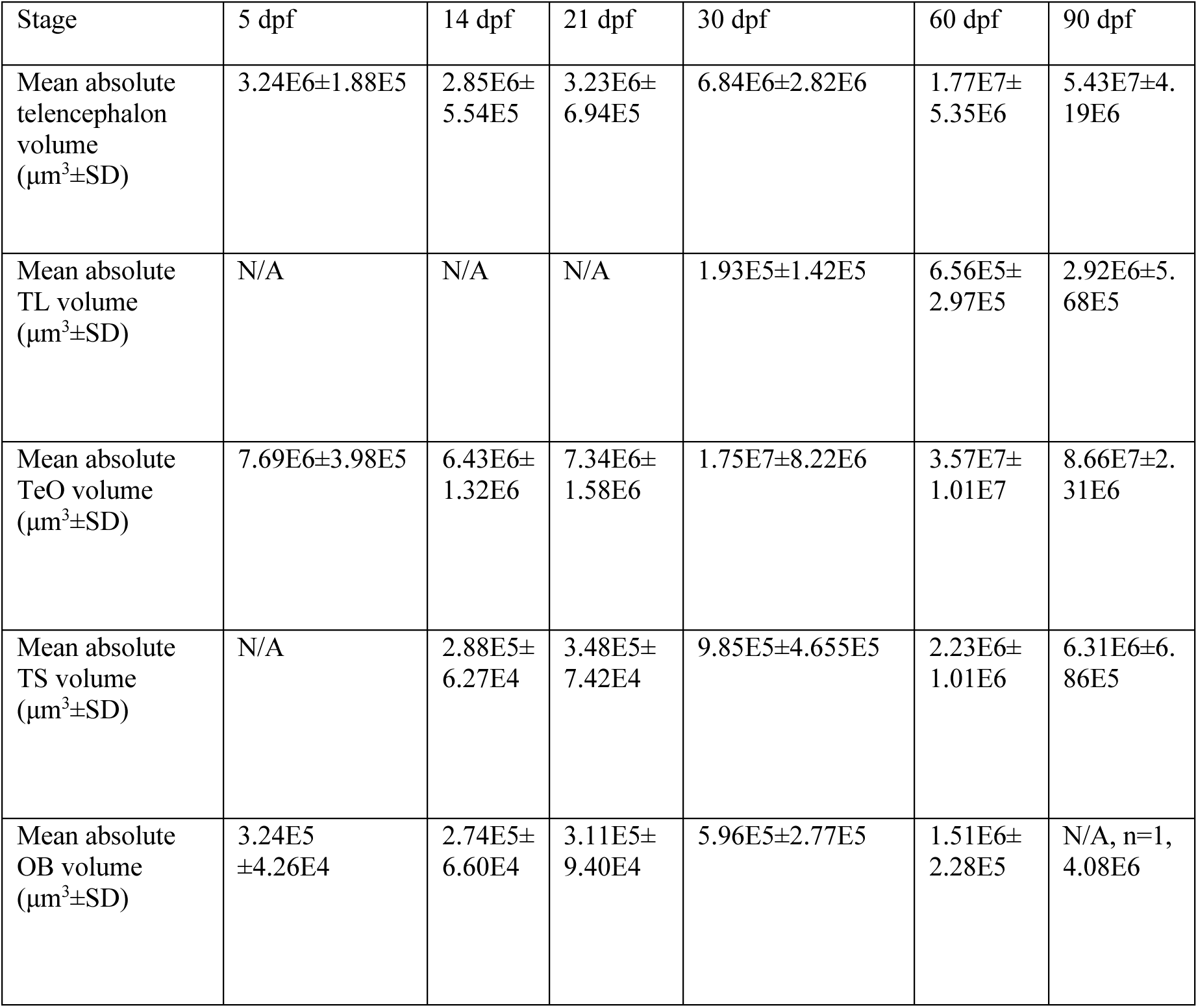
Absolute mean volume (μm^3^±SD) of telencephalon, optic tectum (TeO), torus semicircularis (TS), torus longitudinalis (TL), and olfactory bulb (OB) for each age group. N = same as animals imaged (Table 1) except for 5 dpf TS and 5-21 dpf TL measurements which could not be defined, and 90 dpf OB measurements (see Materials and Methods).

To account for size-related variation in brain region volumes, we standardized each region by calculating normalized gross-brain percentages for each volume. Although the magnitude of variation differed several fold across the brain regions examined (note differences in y-axis), there were significant effects of age for all brain regions (one-way ANOVAs; telencephalon: p<0.001, TL: p<0.001, TeO: p<0.001, TS: p = 0.003, OB: p = 0.02). Post-hoc tests revealed that the mean percentage of the telencephalon in 90 dpf animals was significantly greater than in all other age groups (Fig. 4A; Tables 4, 5). Conversely, the mean percentage of the telencephalon in 30 dpf animals was significantly lower than in 14, 60, and 90 dpf animals, but not significantly different from 5 and 21 dpf animals (Fig. 4A; Tables 4, 5). No significant differences were observed in the mean percentage of the telencephalon among the younger ages (5-21 dpf) when compared to each other or to 60 dpf animals (Fig. 4A; Tables 4, 5). These results indicate that the relative proportion of the telencephalon within the rest of the brain increased after 30 dpf in 60 dpf and 90 dpf animals. A similar pattern, although far less in magnitude, was observed for TL. The TL proportion at 90 dpf was significantly higher than at 30 and 60 dpf animals, and at 60 dpf showed a non-significant trend towards being greater than at 30 dpf (Fig. 4B; Tables 4, 5). The TL could not be differentiated from the midbrain using THG in animals aged 5-21 dpf (see Materials and Methods).

**Figure 4.**
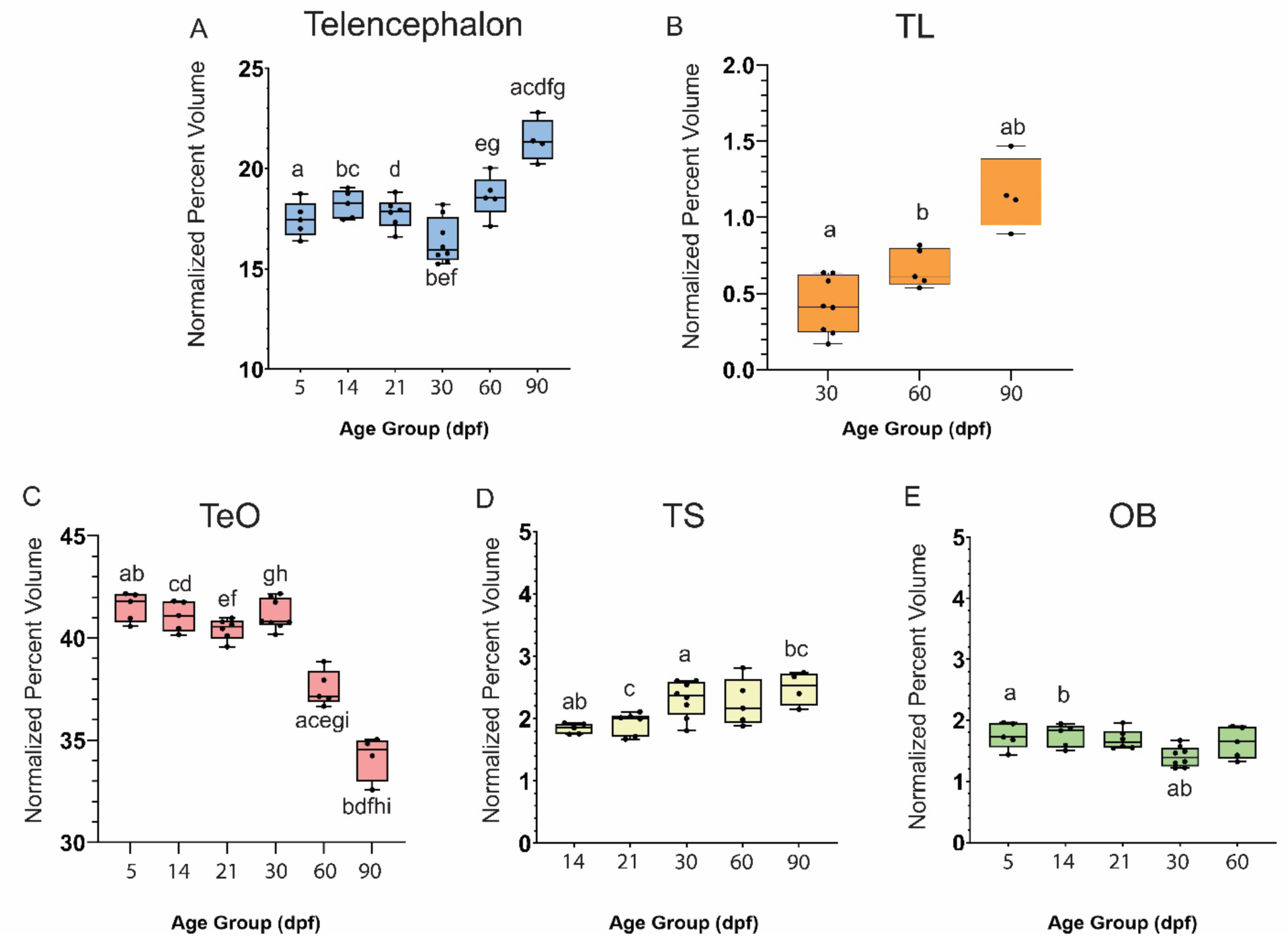
Gross-brain proportion of brain regions across developmental stages shown as boxplots for normalized percent volumes by age group for telencephalon (A), torus longitudinalis (TL, B), optic tectum (TeO, C), torus semicircularis (TS, D) and olfactory bulb (OB, E). Statistically significant differences between two groups are indicated by placing the same letter atop their bars (p < 0.05). For example: the normalized percent volume of the telencephalon of 5 dpf animals is significantly less than that of 90 dpf animals. Therefore, both bars for 5 dpf and 90 dpf animals have the letter “a” above them to signify the significant difference between these age groups for this brain region.

**Table 4.** Mean normalized percent volume (%±SD) of telencephalon, optic tectum (TeO), torus semicircularis (TS), torus longitudinalis (TL), and olfactory bulb (OB) for each age group. N = same as Table 1, except for 5 days post fertilization (dpf) TS, 5-21dpf TL, and 90 dpf OB measures (see Materials and Methods).

| Stage | 5 dpf | 14 dpf | 21 dpf | 30 dpf | 60 dpf | 90 dpf |
| --- | --- | --- | --- | --- | --- | --- |
| Mean normalized percent telencephalon volume (%±SD) | 17.48±0.89 | 18.21±0.70 | 17.77±0.75 | 16.37±1.13 | 18.62±1.04 | 21.41±1.06 |
| Mean normalized percent TL volume (%±SD) | N/A | N/A | N/A | 0.41±0.18 | 0.66±0.12 | 1.15±0.24 |
| Mean normalized percent TeO volume (%±SD) | 41.52±0.72 | 41.06±0.74 | 40.43±0.52 | 41.14±0.74 | 37.53±0.87 | 34.17±1.11 |
| Mean normalized percent TS volume (%±SD) | N/A | 1.84±0.08 | 1.92±0.18 | 2.31±0.29 | 2.26±0.38 | 2.49±0.27 |
| Mean normalized percent OB volume (%±SD) | 1.75±0.22 | 1.75±0.19 | 1.68±0.16 | 1.41±0.17 | 1.64±0.26 | N/A, n=1, 1.63 |

**Table 5:** Post-hoc Tukey HSD Test Results for One-Way ANOVA comparisons on effect of age group (days post fertilization, dpf) on mean normalized percent volume, for each brain region: telencephalon, optic tectum (TeO), torus semicircularis (TS), torus longitudinalis (TL), and olfactory bulb (OB).

| **Telencephalon** | 5 dpf | 14 dpf | 21 dpf | 30 dpf | 60 dpf | 90 dpf |
| --- | --- | --- | --- | --- | --- | --- |
| 5 dpf | - | p=0.825 | p=0.10 | p=0.35 | p=0.431 | p<0.001 |
| 14 dpf |  | - | p=0.972 | p=0.024 | p=0.984 | p<0.001 |
| 21 dpf |  |  | - | p=0.104 | p=0.690 | p<0.001 |
| 30 dpf |  |  |  | - | p=0.004 | p<0.001 |
| 60 dpf |  |  |  |  | - | p=0.002 |
| 90 dpf |  |  |  |  |  | - |
| **TL** |  |  |  |  |  |  |
| 30 dpf |  |  |  | - | p=0.08 | p<0.001 |
| 60 dpf |  |  |  |  | - | p=0.003 |
| **TeO** |  |  |  |  |  |  |
| 5 dpf | - | p=0.932 | p=0.218 | p=0.952 | p<0.001 | p<0.001 |
| 14 dpf |  | - | p=0.757 | p=0.10 | p<0.001 | p<0.001 |
| 21 dpf |  |  | - | p=0.544 | p<0.001 | p<0.001 |
| 30 dpf |  |  |  | - | p<0.001 | p<0.001 |
| 60 dpf |  |  |  |  | - | p<0.001 |
| **TS** |  |  |  |  |  |  |
| 14 dpf |  | - | p=0.98 | p=0.03 | p=0.12 | p=0.009 |
| 21 dpf |  |  | - | p=0.07 | p=0.24 | p=0.02 |
| 30 dpf |  |  |  | - | p=0.1 | p=0.81 |
| 60 dpf |  |  |  |  | - | p=0.69 |
| **OB** |  |  |  |  |  |  |
| 5 dpf | - | p=1 | p=0.98 | p=0.04 | p=0.89 | N/A |
| 14 dpf |  | - | p=0.98 | p=0.04 | p=0.90 | N/A |
| 21 dpf |  |  | - | p=0.1 | p=0.99 | N/A |
| 30 dpf |  |  |  | - | p=0.27 | N/A |

The TeO showed the most pronounced age-related changes in relative size. In contrast to the telencephalon and TL, the TeO proportion in 60 and 90 dpf animals was significantly smaller than in all other age groups (Fig. 4C; Tables 4, 5). The mean percentage of the TeO did not significantly differ among fish ≤ 30 dpf (Fig. 4C, Tables 4, 5).

For the two other modality-dominant brain regions examined, TS and OB, shifts in the relative proportion of each region were far less dramatic compared to the TeO (note differences in y-axis). The TS increased significantly at 30 compared to 14 dpf, and showed a non-significant trend compared to 21 dpf, before remaining consistent in 60 and 90 dpf animals (Fig. 4D; Tables 4, 5). Therefore, the TS displayed a consistent increased proportion at ≥ 30 dpf. The relative proportion of the OB was relatively consistent across all age groups, except for a decrease restricted to 30 dpf (Fig. 4E; Tables 4, 5).

### Changes in Absolute Brain Region Volume

Given the shift after 30 dpf in the relative volume of the different brain regions, we used linear mixed models to examine the effect of brain region and body length on absolute brain volumes for two pooled age groups: animals aged 30 dpf and younger, and older animals of 60 and 90 dpf. Using coefficients from the models, we determined slopes to quantify the relationship between body length and brain region volume. We found that both the telencephalon and TL increase their slopes after 30 dpf, in line with relative proportion findings (Table 6, Fig. 5A). In contrast, the TeO decreases in animals older than 30 dpf, while the slopes for TS and OB are very similar for animals ≤ 30 dpf versus ones that are ≥ 60 dpf. Linear mixed models (Table 7) also revealed significant interactions between body length and brain region affecting raw brain volume, indicating that age-related changes in raw brain volume cannot be attributed solely to increased body length. This likely stems from the notable size differences of these measured brain regions (TeO > telencephalon > TS > OB > TL), which are consistent with patterns observed in other fish species (Wullimann, Rupp, and Reichert, 1996; Nieuwenhuys, ten Donkelaar, and Nicholson, 1998).

**Figure 5:**
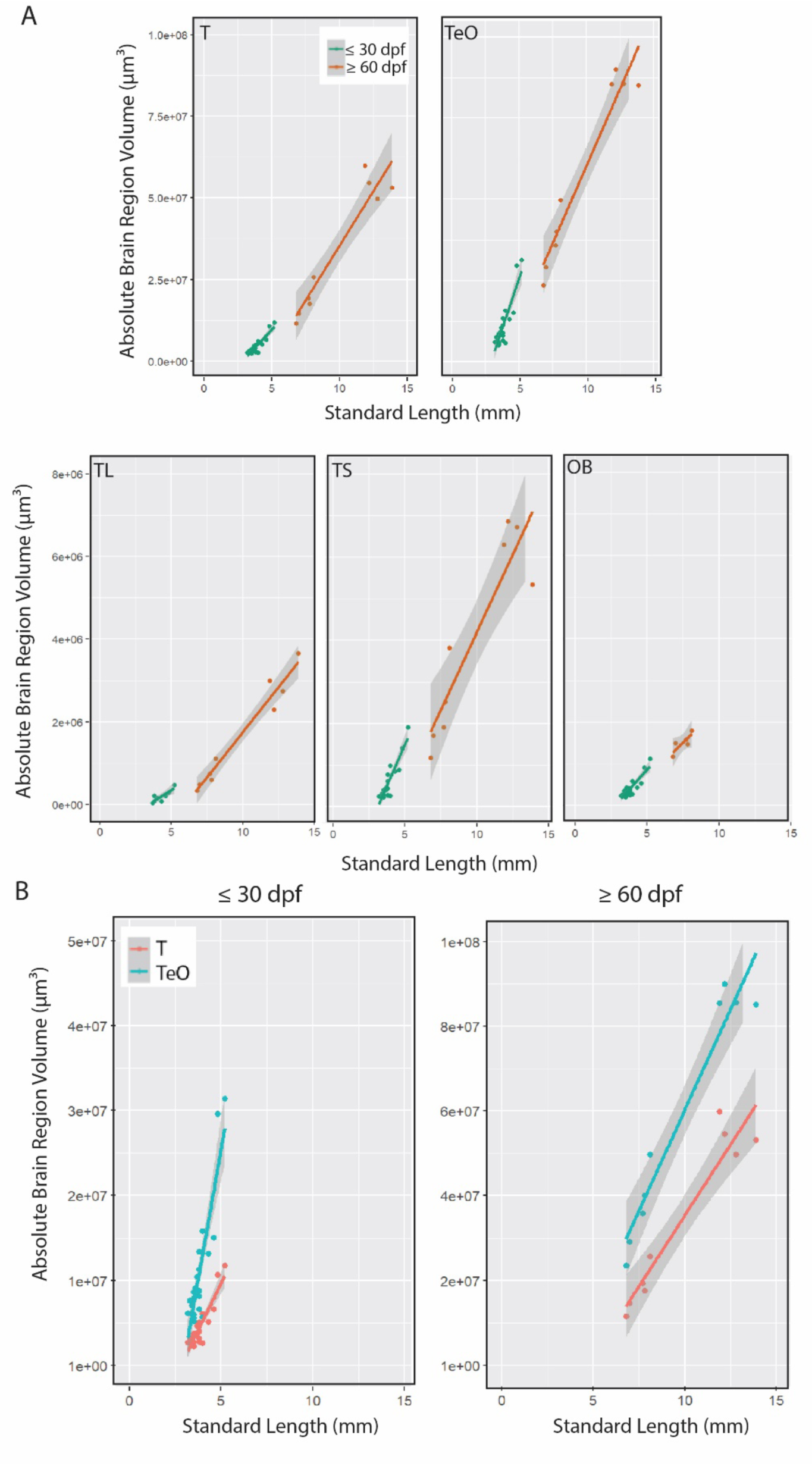
Changes in absolute brain region volume. A. Scatterplots depicting SL (mm) by brain region absolute size (μm^3^). Legend indicates that points and lines indicating values for animals aged 30 days post fertilization (dpf) and younger are in jade green and animals aged older than 30 dpf are in orange. The lines indicate the derived slopes from our linear mixed models. There are five plots for each of the brain regions: telencephalon, optic tectum (TeO), torus semicircularis (TS), torus longitudinalis (TL), and olfactory bulb (OB). B. Scatterplots depicting SL (mm) by brain region absolute size (μm^3^) of T and TeO on same graph, with the first panel demonstrating the derived slope comparisons for animals 30 dpf and younger and the second panel for the derived slope comparisons for animals older than 30 dpf. Legend indicates that points and lines indicating values T values are in coral red and TeO values are in teal. The lines indicate the derived slopes from our linear mixed models.

**Table 6:** Fit slopes for raw volume by standard length in ≤ 30 days post fertilization (dpf) and ≥ 60 dpf fish for telencephalon, optic tectum (TeO), torus semicircularis (TS), torus longitudinalis (TL), and olfactory bulb (OB).

| Brain Region | Slope before and at 30 dpf | Slope after 30 dpf |
| --- | --- | --- |
| Telencephalon | 4.418E6 | 6.69E6 |
| TL | 2.2E5 | 4.36E5 |
| TeO | 1.2301E7 | 9.503E6 |
| TS | 7.93E5 | 7.47E5 |
| OB | 3.83E5 | 3.42E5 |

**Table 7:** Estimated slope coefficients of interactions of body length and brain region on raw brain volume and p-values, comparing telencephalon (T) with optic tectum (TeO), torus semicircularis (TS), torus longitudinalis (TL), and olfactory bulb (OB).

|  | T:Teo | T:TS | T:TL | T:OB |
| --- | --- | --- | --- | --- |
| Age ≤ 30 dpf | Estimated slope: 7.8E6; $p < 0.001$ | Estimated slope: 3.9E6; $p < 0.001$ | Estimated slope: 4.5E6; $p < 0.001$ | Estimated slope: 4.0E6; $p < 0.001$ |
| Age $>$ 30 dpf | Estimated slope: 2.8E6; $p < 0.001$ | Estimated slope: 5.9E6; $p < 0.001$ | Estimated slope: 6.3E6; $p < 0.001$ | Estimated slope: 8.6E6; $p < 0.001$ |

The effect of brain region on raw brain volume with increased body length is 7.8E6 times greater in TeO compared to the telencephalon in animals ≤ 30 dpf, but is decreased to only 2.8E6 times greater in older animals (Table 7; visualization in Fig. 5B). These findings align with our complementary analysis, demonstrating distinct shifts in mean relative percentage for telencephalon and TeO in animals aged 60 and 90 dpf. Comparing the raw brain volume of brain regions smaller than the telencephalon (TL, TS, and OB) shows that the interaction effect of brain region on raw brain volume with increased body length is consistently greater in the telencephalon than in these brain regions in animals ≤ 30 dpf, and then even greater in older animals (Table 7). This finding likely stems from the large shift in size of the telencephalon in older ages compared to the less pronounced increases in the other brain regions.

## DISCUSSION

We used THG microscopy to non-invasively image intact *D. dracula* brains *in vivo* across developmental periods spanning from larval and juvenile stages to adulthood. Measures of relative brain volume and the slope of absolute volume allowed us to characterize developmental shifts in the relative size of five brain regions implicated in higher order (telencephalon, TL) or modality-dominant (TeO, TS, OB) sensory integration. Consistent with our hypothesis, we observed significant changes as larvae became juveniles, where our studied brain regions exhibited allometric growth compared to the rest of the brain. The telencephalon and TeO exhibited the most dramatic changes during the 60-90 dpf period when juveniles transform into adults. The telencephalon showed positive allometry, while the TeO showed negative allometry during this period. Changes for the TL, TS, and OB were quite modest by comparison. Differential investment though was not only evident in the predicted period of 60-90 dpf; there were significant changes in the relative volume of the telencephalon, TS and OB also in 30 dpf animals. As discussed more below, the results set the stage for future studies testing the hypothesis that age-related growth of sensory integration centers in *D. dracula* relate to changes in physiological and behavioral mechanisms from larval to adult life stages.

### Delayed Shifts in Growth of Higher Order Sensory Integration Regions

In terms of relative brain percentage, the brain regions associated with higher order sensory integration, the telencephalon and TL, increased significantly in size starting at 60 dpf, with peak changes at 90 dpf. These shifts correspond with increasing body size and the development of secondary sex characteristics, including the hypertrophied jaw and fangs in adult males. More broadly, this coincides with increased complexity in the behavioral repertoire, as *D. dracula* adults exhibit behaviors not observed in juvenile fish, such as courtship, mating, and reproductive-related aggression (Schulze et al., 2018; Tatarsky et al., 2022). The performance of these adult behaviors may rely, in part, on increased inputs to the telencephalon (e.g., see Northcutt, 2006; Yamamoto and Ito, 2008; Forlano and Bass, 2011) carrying, for example, acoustic and visual information resulting from conspecific sound production and postural displays (see Tatarsky et al., 2022 for *D. dracula*). The TL, a higher order visual area linked to binocular integration, has been suggested to enhance perception in dynamic visual settings (Tesmer et al., 2022) via input from visual and cerebellum-related nuclei in the pretectum (Folgueira et al., 2020). Consequently, the observed volumetric increases in the telencephalon and TL could reflect maturation-related changes in these and other sensory-related mechanisms.

*Danionella* species may offer unique opportunities to establish the interplay between intrinsic and extrinsic factors contributing to the differential development of sensory systems because of the non-terminal nature of neuroimaging methods such as THG that allow for the re-imaging of individual fish as they age (Lam, 2022; Akbari et al., 2022; Rajan et al., 2022; Akbari et al., 2024). For example, are especially prominent increases in telencephalic size primarily due to uniform growth across the entire region or the growth of specific subregions, such as the lateral pallium (Dirian et al., 2014) or the discrete lobes recognized in the area dorsalis of *D. dracula* (Fig. 3B)? How are the equally striking decreases in TeO size observed over a similar timeframe related to changing visual and non-visual processing, related in part to motor function and behavioral output (e.g., see Thompson et al., 2016; Pietri et al., 2017)? More specifically, such a detailed analysis could uncover insights into modifications in sensation related to spatial navigation and memory in *Danionella* and other fish species, and vertebrates in general (Rodríguez et al., 2002; Portavella & Vargas, 2005; Vargas et al., 2006; Yamamoto et al., 2007; Nieuwenhuys, 2011; Giassi et al., 2012; Costa et al., 2011; Rajan et al., 2022; Lee & Briggman, 2023; Zada et al., 2024).

### Sensory Brain Region Investment over Development

As *D. dracula* age, the TeO, OB, and TS all increase in absolute size, but their proportional changes in relation to brain size differ significantly. While the TeO’s proportion decreases in animals older than 30 dpf, the proportions of the OB and TS remain relatively consistent. This may be due to the larval brain of cyprinids often exhibiting large optic lobes (Brandstatter and Kotrschal, 1989), along with early development of the eye and retinal connections with the tectum (Kawamura et al., 1984; Stuermer and Raymond, 1989). Researchers have observed a relatively large TeO in fish larvae, with its relative size decreasing over the course of development, in many species of fish, including salmoniformes, the Antarctic silverfish, the deep-sea grenadier, elasmobranchs, weakly electric gymnotid fish and cyprinids such as roach, bream, common carp, and sabre carp (Cadwallader, 1975; Brandstatter and Kotrschal, 1989, 1990; Montgomery et al., 1997; Wagner, 2003; Lisney et al., 2007; Iribarne and Castello, 2014). This decrease in the relative size of TeO over development could stem from initial size constraints in larvae and subsequent release of these constraints during growth (Brandstätter and Kotrschal, 1990). Additionally, it may reflect shifts in sensory capabilities underlying changes in early feeding behaviors or visual processing in later stages (Toyoda and Uematsu, 1994; Iribarne and Castello, 2014). For example, many fish larvae feed during the day in relatively high light levels (Sandy and Blaxter, 1980), but juveniles and adults may shift towards dawn, dusk, and even nocturnal activity (Brandstätter and Kotrschal, 1990). Regarding social behavior, during development up to 30 dpf, zebrafish begin to demonstrate visually-guided aspects of complex behaviors including social attraction, shoaling, and social cueing (Dreosti et al., 2015; Gerlai, 2017; Larsch and Baier, 2018; Stednitz et al., 2018; Kappel et al., 2022). Recent work in *D. cerebrum* as well, a sister species of *D. dracula*, has indicated that schooling develops sequentially, starting with social avoidance at 14 dpf, aggregation at 30 dpf, and postural alignment and coordination between 40 and 60 dpf (Zada et al., 2024). Our results could indicate increased reliance in visual processing before 30 dpf in *D. dracula* juveniles compared to other senses.

As noted above, while various species of cyprinids have highly similar brain morphologies in the larval and early-juvenile stages, for example large optic lobes, species-specific brain structures can emerge prominently during the late-larval and juvenile periods (Brandstätter and Kotrschal, 1989, 1990). For example, in weakly electric fish, the marked increase in the relative size of the electrosensory lateral line lobe (ELL) from larval to juvenile stages, alongside the reduction in the relative size of the TeO, suggests age-related tradeoffs in the relative importance of visual and electrosensory information, likely associated with shifts in feeding behavior (Iribarne and Castello, 2014). In contrast to the TeO, our OB results indicate a consistent investment in olfaction over development, in line with early development of olfactory processing and discrimination in other fish species. For example, zebrafish larvae can detect amino acid odorants and show aversive responses to cysteine shortly after hatching (Lindsay and Vogt, 2004; Vitebsky et al., 2005; Miyasaka et al., 2013). However, zebrafish only show behavioral responses to alarm substance and pheromones involved in mating later in life (Waldman, 1982; Darrow and Harris, 2004). Perhaps *D. dracula* develops olfactory complexity more slowly and consistently over development compared to visual complexity mediated by the TeO.

We were originally curious if there may be a significant critical period for development of the auditory and lateral line recipient TS and thus increased size in that brain region, given that *D. dracula* males produce sounds robustly during aggressive interactions in community settings (Tatarsky et al., 2022) and exhibit accelerated formation of the Weberian apparatus, a chain of ossicles linking the swim bladder to the inner ear that enhances the sense of hearing (Braun and Grande, 2008; Conway, Kubicek and Britz, 2021). We found that TS was relatively consistent in investment over development but did display an increased brain proportion between 21-30 dpf that is maintained into adulthood, i.e., 90 dpf. We could not distinguish between auditory and lateral line divisions of the TS, although the growth spurt coincides with the time period over which the Weberian apparatus fully develops (Conway, Kubicek and Britz, 2021). The exact onset of sound production in *D. dracula* is not yet known, but it may be related to the increased separation of the two chambers of the swim bladder that we observed from 14-30 dpf (Fig. 1D). The anterior portion of the swim bladder alone is associated with the Weberian apparatus, as well as a sonic “drumming apparatus” associated with the swim bladder in adult males, but not females (Britz and Conway, 2016; Cook et al., 2024). In *D. cerebrum*, males begin to produce sounds in between 6.5 and 8 weeks of development, concurrent with development of the sound-producing drumming apparatus (Groneberg et al., 2024). Future research into hearing and acoustic communication should assess auditory and lateral line sensitivity and acoustic response behavior at different ages.

To build upon these initial findings on contrasting patterns of sensory brain region development, it would be important to assess each region’s respective sensory integration capacity aligned with changes in the performance of pre-adult and adult social behaviors. This approach aligns with the strategies being developed in zebrafish research (Stednitz et al., 2018; Kappel et al., 2022), but with the addition of a robustly measurable acoustic component (see Tatarsky et al., 2022).

### Concluding comments

We demonstrate that *D. dracula,* a miniature transparent species amenable to whole-brain neuroimaging using THG, exhibits changes in the relative volume of sensory-related regions from larval to adult stages of development. A single timepoint, 30 dpf, stands out as a developmental milestone for significant shifts in the relative volume of the different brain regions examined here. Peak increases witnessed for the telencephalon and TL coincide with the onset of adulthood and complex social behaviors related to reproduction, namely courtship and mating. The especially large investment in the TeO at ≤ 30 dpf and relatively stable investment in the olfactory-recipient OB throughout development stand out, highlighting their importance to the performance of visual- and olfactory-dependent behaviors at these stages, all of which remains to be investigated. From a broader functional perspective, increased TS investment at 30 dpf overlaps the timepoint when the swim bladder has split into separate anterior and posterior divisions, which may further coincide with the onset of sound production.

Third harmonic generation microscopy could further be used in *D. dracula* to image and compare male and female brains, especially given the robust differences in sonic and jaw extension behaviors between the sexes (Tatarsky et al., 2022). Species differences can also be readily assessed, such as comparing the brain of *D. dracula* with that of its more highly sonic sister species, *D. cerebrum* who lack the jaw extension postural display (Schulze et al., 2018; Akbari et al., 2024). Finally, the application of THG in these and other small transparent species allows investigations of the neural basis of behavior using non-terminal whole-brain neuroimaging methods and possible chronic imaging of the same animals as they develop into adulthood, in combination with any genetic tools that are actively being created within this genus (Schulze et al., 2018; Akbari et al., 2022; Akbari et al., 2024).

## Acknowledgements

We thank Jonathan Perelmuter and Sarah Campbell for many helpful comments on the manuscript, Joe Fetcho for research guidance and use of IMARIS software, and Stephen Parry at the Cornell Statistical Consulting Unit for discussions on data analysis.

## REFERENCES

Ahn, S. J., Ruiz-Uribe, N. E., Li, B., Porter, J., Sakadzic, S., & Schaffer, C. B. (2020). Label-free assessment of hemodynamics in individual cortical brain vessels using third harmonic generation microscopy. Biomedical optics express, 11(5), 2665–2678.

Akbari, N., Tatarsky, R. L., Kolkman, K. E., Fetcho, J. R., Bass, A. H., & Xu, C. (2022). Whole-brain optical access in a small adult vertebrate with two-and three-photon microscopy. Iscience, 25(10).

Akbari, N., Tatarsky, R. L., Kolkman, K. E., Fetcho, J. R., Xu, C., & Bass, A. H. (2024). Label-free, whole-brain in vivo mapping in an adult vertebrate with third harmonic generation microscopy. Journal of Comparative Neurology, 532(4), e25614.

Axelrod, C. J., Laberge, F., & Robinson, B. W. (2018). Intraspecific brain size variation between coexisting sunfish ecotypes. Proceedings of the Royal Society B, 285(1890), 20181971.

Barton, R., Purvis, A., & Harvey, P. H. (1995). Evolutionary radiation of visual and olfactory brain systems in primates, bats and insectivores. Philosophical Transactions of the Royal Society of London. Series B: Biological Sciences, 348(1326), 381–392.

Barton, R. A., & Harvey, P. H. (2000). Mosaic evolution of brain structure in mammals. Nature, 405(6790), 1055–1058.

Bitplane (2018) Imaris 9.2 Reference Manual. Oxford Instruments, Abingdon, United Kingdom.

Brandstätter, R., & Kotrschal, K. (1989). Life History of Roach, Rutilus rutilus (Cyprinidae, Teleostei) A Qualitative and Quantitative Study on the Development of Sensory Brain Areas. Brain, behavior and evolution, 34(1), 35–42.

Brandstätter, R., & Kotrschal, K. (1990). Brain growth patterns in four European cyprinid fish species (Cyprinidae, Teleostei): roach (Rutilus rutilus), bream (Abramis brama), common carp (Cyprinus carpio) and sabre carp (Pelecus cultratus). Brain, behavior and evolution, 35(4), 195–211.

Braun, C. B., & Grande, T. (2008). Evolution of peripheral mechanisms for the enhancement of sound reception. Fish Bioacoustics: With 81 Illustrations, 99–144.

Britz, R., Conway, K. W., & Rüber, L. (2009). Spectacular morphological novelty in a miniature cyprinid fish, Danionella dracula n. sp. Proceedings of the Royal Society B: Biological Sciences, 276(1665), 2179–2186.

Britz, R., & Conway, K. W. (2016). Danionella dracula, an escape from the cypriniform Bauplan via developmental truncation?. Journal of morphology, 277(2), 147–166.

Britz, R., Conway, K. W., & Rüber, L. (2021). The emerging vertebrate model species for neurophysiological studies is Danionella cerebrum, new species (Teleostei: Cyprinidae). Scientific reports, 11(1), 18942.

Cadwallader, P. L. (1975). Relationship between brain morphology and ecology in New Zealand Galaxiidae, particularly Galaxias vulgaris (Pisces: Salmoniformes). New Zealand journal of zoology, 2(1), 35–43.

Chow, D. M., Sinefeld, D., Kolkman, K. E., Ouzounov, D. G., Akbari, N., Tatarsky, R., … & Fetcho, J. R. (2020). Deep three-photon imaging of the brain in intact adult zebrafish. Nature Methods, 17(6), 605–608.

Conway, K. W., Kubicek, K.M., & Britz, R. (2021). Extreme evolutionary shifts in developmental timing establish the miniature Danionella as a novel model in the neurosciences. Developmental Dynamics, 250(4), 601–611.

Costa, S. S., Andrade, R., Carneiro, L. A., Gonçalves, E. J., Kotrschal, K., & Oliveira, R. F. (2011). Sex differences in the dorsolateral telencephalon correlate with home range size in blenniid fish. Brain, Behavior and Evolution, 77(1), 55–64.

Darrow, K. O., & Harris, W. A. (2004). Characterization and development of courtship in zebrafish, Danio rerio. Zebrafish, 1(1), 40–45.

Davis, R. E., & Northcutt, R. G. (Eds.). (1983). Fish Neurobiology: Higher Brain Areas and Functions. University of Michigan Press.

DeMarco, E., Tesmer, A. L., Kawakami, K., & Robles, E. (2021). Pyramidal neurons of the zebrafish tectum receive highly convergent input from torus longitudinalis. Frontiers in Neuroanatomy, 15, 636683.

Dirian, L., Galant, S., Coolen, M., Chen, W., Bedu, S., Houart, C., … & Foucher, I. (2014). Spatial regionalization and heterochrony in the formation of adult pallial neural stem cells. Developmental cell, 30(2), 123–136.

Dreosti, E., Lopes, G., Kampff, A. R., & Wilson, S. W. (2015). Development of social behavior in young zebrafish. Frontiers in neural circuits, 9, 39.

Folgueira, M., Anadón, R., & Yáñez, J. (2004). Experimental study of the connections of the telencephalon in the rainbow trout (Oncorhynchus mykiss). II: Dorsal area and preoptic region. Journal of Comparative Neurology, 480(2), 204–233.

Forlano, P. M., & Bass, A. H. (2011). Neural and hormonal mechanisms of reproductive-related arousal in fishes. Hormones and behavior, 59(5), 616–629.

Gerlai, R. (2017). Zebrafish and relational memory: Could a simple fish be useful for the analysis of biological mechanisms of complex vertebrate learning?. Behavioural Processes, 141, 242–250.

Giassi, A. C., Duarte, T. T., Ellis, W., & Maler, L. (2012). Organization of the gymnotiform fish pallium in relation to learning and memory: II. Extrinsic connections. Journal of Comparative Neurology, 520(15), 3338–3368.

Gonda, A., Herczeg, G., & Merilä, J. (2009). Habitat-dependent and-independent plastic responses to social environment in the nine-spined stickleback (Pungitius pungitius) brain. Proceedings of the Royal Society B: Biological Sciences, 276(1664), 2085–2092.

Goode, C., Voeun, M., Ncube, D., Eisen, J., Washbourne, P., & Tallafuss, A. (2021). Late onset of Synaptotagmin 2a expression at synapses relevant to social behavior. Journal of Comparative Neurology, 529(9), 2176–2188.

Groneberg, A. H., Dressler, L. E., Kadobianskyi, M., Muller, J., & Judkewitz, B. (2024). Development of sound production in Danionella cerebrum. bioRxiv, 2024-03.

Farrar, M. J., Wise, F. W., Fetcho, J. R., & Schaffer, C. B. (2011). In vivo imaging of myelin in the vertebrate central nervous system using third harmonic generation microscopy. Biophysical journal, 100(5), 1362–1371.

Hinaux, H., Devos, L., Blin, M., Elipot, Y., Bibliowicz, J., Alié, A., & Rétaux, S. (2016). Sensory evolution in blind cavefish is driven by early embryonic events during gastrulation and neurulation. Development, 143(23), 4521–4532.

Huber, R., van Staaden, M. J., Kaufman, L. S., & Liem, K. F. (1997). Microhabitat use, trophic patterns, and the evolution of brain structure in African cichlids. Brain, Behavior and Evolution, 50(3), 167–182.

Iribarne, L., & Castelló, M. E. (2014). Postnatal brain development of the pulse type, weakly electric gymnotid fish Gymnotus omarorum. Journal of Physiology-Paris, 108(2-3), 47–60.

Jaggard, J. B., Lloyd, E., Yuiska, A., Patch, A., Fily, Y., Kowalko, J. E., … & Keene, A. C. (2020). Cavefish brain atlases reveal functional and anatomical convergence across independently evolved populations. Science advances, 6(38), eaba3126.

Kappel, J. M., Förster, D., Slangewal, K., Shainer, I., Svara, F., Donovan, J. C., … & Larsch, J. (2022). Visual recognition of social signals by a tectothalamic neural circuit. Nature, 608(7921), 146–152.

Kawamura, G., Tsuda, R., Kumai, H., & Ohashi, S. (1984). The visual cell morphology of Pagrus major and its adaptive changes with shift from pelagic to benthic habitats. Nippon Suisan Gakkaishi, 50, 1975–1980.

Keesey, I. W., Grabe, V., Knaden, M., & Hansson, B. S. (2020). Divergent sensory investment mirrors potential speciation via niche partitioning across Drosophila. Elife, 9, e57008.

Kotrschal, A., Rogell, B., Bundsen, A., Svensson, B., Zajitschek, S., Brännström, I., … & Kolm, N. (2013). Artificial selection on relative brain size in the guppy reveals costs and benefits of evolving a larger brain. Current Biology, 23(2), 168–171.

Kunst, M., Laurell, E., Mokayes, N., Kramer, A., Kubo, F., Fernandes, A. M., … & Baier, H. (2019). A cellular-resolution atlas of the larval zebrafish brain. Neuron, 103(1), 21–38.

Lam, P. Y. (2022). Longitudinal in vivo imaging of adult Danionella cerebrum using standard confocal microscopy. Disease Models & Mechanisms, 15(12), dmm049753.

Larsch, J., & Baier, H. (2018). Biological motion as an innate perceptual mechanism driving social affiliation. Current Biology, 28(22), 3523–3532.

Lee, T. J., & Briggman, K. L. (2023). Visually guided and context-dependent spatial navigation in the translucent fish Danionella cerebrum. Current Biology, 33(24), 5467–5477.

Lenth, R.V. (2016). Least-Squares Means: The R Package lsmeans. Journal of Statistical Software, 69(1), 1–33.

Lindsay, Sara M., and Richard G. Vogt. “Behavioral responses of newly hatched zebrafish (Danio rerio) to amino acid chemostimulants.” Chemical Senses 29.2 (2004): 93–100.

Lisney, T. J., Bennett, M. B., & Collin, S. P. (2007). Volumetric analysis of sensory brain areas indicates ontogenetic shifts in the relative importance of sensory systems in elasmobranchs. Raffles Bulletin of Zoology, 55(Supp. 14), 7–15.

Lopes Corrêa, S. A., Grant, K., & Hoffmann, A. (1998). Afferent and efferent connections of the dorsocentral telencephalon in an electrosensory teleost, Gymnotus carapo. Brain behavior and evolution, 52(2), 81–98.

McArthur, K. L., Chow, D. M., & Fetcho, J. R. (2020). Zebrafish as a model for revealing the neuronal basis of behavior. In The zebrafish in biomedical research (pp. 593–617). Academic Press.

Miyasaka, N., Wanner, A. A., Li, J., Mack-Bucher, J., Genoud, C., Yoshihara, Y., & Friedrich, R. W. (2013). Functional development of the olfactory system in zebrafish. Mechanisms of development, 130(6-8), 336–346.

Montgomery, J. C., & Sutherland, K. B. W. (1997). Sensory development of the Antarctic silverfish Pleuragramma antarcticum: a test for the ontogenetic shift hypothesis. Polar Biology, 18, 112–115.

Moran, D., Softley, R., & Warrant, E. J. (2015). The energetic cost of vision and the evolution of eyeless Mexican cavefish. Science advances, 1(8), e1500363.

Niven, J. E., & Laughlin, S. B. (2008). Energy limitation as a selective pressure on the evolution of sensory systems. Journal of Experimental Biology, 211(11), 1792–1804.

Nieuwenhuys, R. (2011). The development and general morphology of the telencephalon of actinopterygian fishes: synopsis, documentation and commentary. Brain Structure and Function, 215, 141–157.

Nieuwenhuys, R., Ten Donkelaar, H. J., & Nicholson, C. (1998). The central nervous system of vertebrates. Berlin, Germany: Springer.

Nixdorf-Bergweiler, B., & und Halbach, V. V. B. (2004). Major sex differences in the development of myelination are prominent in song system nucleus HVC but not lMAN. Animal biology, 54(1), 27–43.

Northcutt, R. G. (2006). Connections of the lateral and medial divisions of the goldfish telencephalic pallium. Journal of Comparative Neurology, 494(6), 903–943.

O’Donnell, S., Clifford, M., & Molina, Y. (2011). Comparative analysis of constraints and caste differences in brain investment among social paper wasps. Proceedings of the National Academy of Sciences, 108(17), 7107–7112.

Pallotto, M., Watkins, P. V., Fubara, B., Singer, J. H., & Briggman, K. L. (2015). Extracellular space preservation aids the connectomic analysis of neural circuits. Elife, 4, e08206.

Pietri, T., Romano, S. A., Pérez-Schuster, V., Boulanger-Weill, J., Candat, V., & Sumbre, G. (2017). The emergence of the spatial structure of tectal spontaneous activity is independent of visual inputs. Cell reports, 19(5), 939–948.

Portavella, M., & Vargas, J. P. (2005). Emotional and spatial learning in goldfish is dependent on different telencephalic pallial systems. European Journal of Neuroscience, 21(10), 2800–2806.

Prechtl, J. C., von der Emde, G., Wolfart, J., Karamürsel, S., Akoev, G. N., Andrianov, Y. N., & Bullock, T. H. (1998). Sensory processing in the pallium of a mormyrid fish. Journal of Neuroscience, 18(18), 7381–7393.

Rajan, G., Lafaye, J., Faini, G., Carbo-Tano, M., Duroure, K., Tanese, D., … & Del Bene, F. (2022). Evolutionary divergence of locomotion in two related vertebrate species. Cell Reports, 38(13).

R Core Team (2021) “R: A language and environment for statistical computing.” Vienna, Austria: R Foundation for Statistical Computing. Available at: https://www.R-project.org/.

Rodríguez, F., López, C., Vargas, J.P., Gómez, Y., Broglio, C., Salas, C. (2002) Conservation of spatial memory function in the pallial forebrain of reptiles and ray-finned fishes. J Neurosci. 22:2894–2903.

Sandy, J. M., & Blaxter, J. H. S. (1980). A study of retinal development in larval herring and sole. Journal of the Marine Biological Association of the United Kingdom, 60(1), 59–71.

Schulze, L., Henninger, J., Kadobianskyi, M., Chaigne, T., Faustino, A. I., Hakiy, N., … & Judkewitz, B. (2018). Transparent Danionella translucida as a genetically tractable vertebrate brain model. Nature methods, 15(11), 977–983.

Sheehan, Z. B., Kamhi, J. F., Seid, M. A., & Narendra, A. (2019). Differential investment in brain regions for a diurnal and nocturnal lifestyle in Australian Myrmecia ants. Journal of Comparative Neurology, 527(7), 1261–1277.

Shinozuka, K., & Watanabe, S. (2004). Effects of telencephalic ablation on shoaling behavior in goldfish. Physiology & behavior, 81(1), 141–148.

Stednitz, S. J., McDermott, E. M., Ncube, D., Tallafuss, A., Eisen, J. S., & Washbourne, P. (2018). Forebrain control of behaviorally driven social orienting in zebrafish. Current biology, 28(15), 2445–2451.

Stöckl, A., Heinze, S., Charalabidis, A., El Jundi, B., Warrant, E., & Kelber, A. (2016). Differential investment in visual and olfactory brain areas reflects behavioural choices in hawk moths. Scientific Reports, 6(1), 26041.

Striedter, G. F. (2005). Principles of brain evolution. Sinauer associates.

Striedter, G. F., & Northcutt, R. G. (2022). The independent evolution of dorsal pallia in multiple vertebrate lineages. Brain Behavior and Evolution, 96(4-6), 200–211.

Stuermer, C. A., & Raymond, P. A. (1989). Developing retinotectal projection in larval goldfish. Journal of Comparative Neurology, 281(4), 630–640.

Tatarsky, R. L., Guo, Z., Campbell, S. C., Kim, H., Fang, W., Perelmuter, J. T., … & Bass, A. H. (2022). Acoustic and postural displays in a miniature and transparent teleost fish, Danionella dracula. Journal of Experimental Biology, 225(16), jeb244585.

Teles, M. C., Almeida, O., Lopes, J. S., & Oliveira, R. F. (2015). Social interactions elicit rapid shifts in functional connectivity in the social decision-making network of zebrafish. Proceedings of the Royal Society B: Biological Sciences, 282(1816), 20151099.

Tesmer, A. L., Fields, N. P., & Robles, E. (2022). Input from torus longitudinalis drives binocularity and spatial summation in zebrafish optic tectum. BMC biology, 20(1), 24.

Thompson, A. W., Vanwalleghem, G. C., Heap, L. A., & Scott, E. K. (2016). Functional profiles of visual-, auditory-, and water flow-responsive neurons in the zebrafish tectum. Current Biology, 26(6), 743–754.

Toyoda, J., & Uematsu, K. (1994). Brain morphogenesis of the red sea bream, Pagrus major (Teleostei). Brain, Behavior and Evolution, 44(6), 324–337.

Ullmann, J. F., Cowin, G., Kurniawan, N. D., & Collin, S. P. (2010). A three-dimensional digital atlas of the zebrafish brain. Neuroimage, 51(1), 76–82.

Vargas, J. P., Bingman, V. P., Portavella, M., & López, J. C. (2006). Telencephalon and geometric space in goldfish. European Journal of Neuroscience, 24(10), 2870–2878.

Vitebsky, A., Reyes, R., Sanderson, M. J., Michel, W. C., & Whitlock, K. E. (2005). Isolation and characterization of the laure olfactory behavioral mutant in the zebrafish, Danio rerio. Developmental dynamics, 234(1), 229–242.

Wagner, H. J. (2003). Volumetric analysis of brain areas indicates a shift in sensory orientation during development in the deep-sea grenadier Coryphaenoides armatus. Marine Biology, 142, 791–797.

Waldman, B. (1982). Quantitative and developmental analyses of the alarm reaction in the zebra danio, Brachydanio rerio. Copeia, 1–9.

Weigelin, B., Bakker, G. J., & Friedl, P. (2016). Third harmonic generation microscopy of cells and tissue organization. Journal of Cell Science, 129(2), 245–255.

Wullimann, M.F., Rupp, B., Reichert, H. (1996) Neuroanatomy of the zebrafish brain: a topological atlas. Birkhäuser, Basel: Springer.

Yamamoto, Y., Stock, D. W., & Jeffery, W. R. (2004). Hedgehog signalling controls eye degeneration in blind cavefish. Nature, 431(7010), 844–847.

Yamamoto, N., Ishikawa, Y., Yoshimoto, M., Xue, H. G., Bahaxar, N., Sawai, N., … & Ito, H. (2007). A new interpretation on the homology of the teleostean telencephalon based on hodology and a new eversion model. Brain, Behavior and Evolution, 69(2), 96–104.

Yamamoto, N., & Ito, H. (2008). Visual, lateral line, and auditory ascending pathways to the dorsal telencephalic area through the rostrolateral region of the lateral preglomerular nucleus in cyprinids. Journal of Comparative Neurology, 508(4), 615–647.

Yamamoto, Y., Byerly, M. S., Jackman, W. R., & Jeffery, W. R. (2009). Pleiotropic functions of embryonic sonic hedgehog expression link jaw and taste bud amplification with eye loss during cavefish evolution. Developmental biology, 330(1), 200–211.

Yi, W., Mueller, T., Rücklin, M., & Richardson, M. K. (2022). Developmental neuroanatomy of the rosy bitterling Rhodeus ocellatus (Teleostei: Cypriniformes)—A microCT study. Journal of Comparative Neurology, 530(12), 2132–2153.

Zada, D., Schulze, L., Yu, J.-H., Tarabishi, P., Napoli, J. L., Milan, J., & Lovett-Barron, M. (2024). Development of neural circuits for social motion perception in schooling fish. Current Biology, 34, 1–123.

